# Captive-breeding and catch-and-release’s effects on the reproductive success of Atlantic salmon (*Salmo salar* L.)

**DOI:** 10.1101/2021.04.06.438651

**Authors:** R. Bouchard, K. Wellband, L. Lecomte, L. Bernatchez, J. April

## Abstract

Captive-breeding programs as well as and catch-and-release are among the most commonly adopted conservation practices in recreational fisheries. However, risks and benefits associated with their implementation are rarely evaluated. In the case of Atlantic Salmon, while previous studies revealed that captive-bred fish show reduced fitness compared to their wild counterparts in nature. Yet, few examined the extent and causes of their reduced reproductive success or directly compared their contribution to enhance genetic diversity to that of wild fish, including mature male parr. Furthermore, only one study specifically measured the reproductive success of caught and released Atlantic salmon in natural settings, and no study to date evaluated if released salmon are able to reproduce when released at temperature above 20°C which is known to increase post-release mortality. Here, we use high-throughput microsatellite sequencing of 38 loci to accurately assign 2500 offspring to a comprehensive set of possible parents from a supplemented Atlantic salmon population in Québec, Canada. The resolved molecular pedigree provided informative insight on the reproductive pattern of both captive-bred salmon and caught-and-released salmon. Captive-bred salmon had fewer partners than their wild conspecifics which lead to a significant reduction of reproductive success relative to that of their wild counterparts. Supplementation of captive-bred salmon significantly contributed to increase genetic diversity but mature male parr did so to an even greater extent and significantly inflated the number of alleles found among offspring. Moreover, our results showed that that at least 83% of caught-and-released salmon did successfully reproduced although caught-and-released female salmon have a significantly reduced reproductive success, averaging 73% of the reproductive output of non-caught salmon. Reproductive success of released salmon was not influenced by water temperature over 20°C which suggests either that the studied population is locally adapted to warm waters or that they behaviorally regulated body temperature by accessing nearby thermal refugia. Our results should help refining managers’ ability to analyze the risks and benefits associated with captive-breeding and catch-and-release, and thus, optimize conservation practices used for the preservation of Atlantic salmon populations.

## Introduction

In industrialized countries from temperate regions of the world, the rise of personal wealth and leisure has generally shifted freshwater subsistence and commercial fisheries towards recreational fisheries (Arlinghaus & Cooke, 2009) with an estimate of at least 220 million recreational fishers worldwide (FAO, 2018). This is more than five times the number of commercial capture fishermen (Arlinghaus et al. 2015). Salmonids rank among the most exploited species by recreational fishery despite numerous populations having experienced worldwide decline over the past century (Post et al. 2002). Since recreational fisheries can have negative impacts on exploited populations through overharvesting (Cooke & Cowx, 2004; Lewin et al. 2006), careful management is needed in order to sustain this important socio-economic sector (Arlinghaus et al. 2019). Two main practices currently exist to mitigate its impact, namely captive-breeding programs for supplementation and catch-and-release fishing.

Captive-breeding programs are widely used for the conservation and restoration of threatened and endangered species with the ultimate goal of maintaining wildlife biodiversity, including genetic diversity and fitness within population (Fraser, 2008). Although this practice has continuously adapted in parallel with increased knowledge, decades of studies have shown that captive-breeding programs may cause a range of negative effects on stocked populations (e.g. Ryman & Laikre, 1991; Hindar et al. 1991; Waples, 1991; Valiquette et al. 2014; Waples et al. 2016; Hagen et al. 2020). For instance, captive rearing has been shown to induce genetic and epigenetic changes which may be due either to unintentional domestication selection or epigenetic reprogramming in hatchery environments (Christie et al. 2012; Le Luyer et al. 2017). Genetic changes result from directional selective pressure in the hatchery which comes at the expense of adaptation to the natural environment (Araki et al. 2008). Consequently, individuals released in the wild have lower fitness compared to naturally produced conspecifics (Araki et al. 2007a). Increased understanding pertaining to (epi)-genetic domestication effects has led to the recommendation that time spent in captivity should be minimized and to use naturally produced fish as broodstock to avoid cumulative fitness decline (Willoughby et Christie, 2019).

Catch-and-release (C&R) angling is a promising complement to captive-breeding programs which is generally applied and promoted to cope with high angling intensity (Arlinghaus et al. 2007). Notably, C&R allows the social and economic benefits pertaining to recreational fishery to persist, even when stock abundance is low (Cooke & Cowk, 2004). Hence, properly applied C&R can provide a long-term management answer to potential angling-induced impacts on fish population unlike captive-breeding. Nonetheless, the success of C&R as a management tool will depend on the ability of the released fish to survive and contribute to reproduction (Arlinghaus et al. 2007; Brownscombe et al. 2017). Indeed, C&R angling can virtually affect or alter almost all biological processes that contribute to survival and an organism’s role in the community (Cooke & Sneddon, 2007). For instance, mortality rates can be highly variable, ranging from 0% to 95% across a variety of marine and freshwater species (Bartholomew & Bohnsack, 2005). While several studies documented survival of C&R fish, few examined their reproductive success (Booth et al. 1995; Whoriskey et al. 2000; Thorstad et al. 2003; Richard et al. 2013). Other than the study by Richard et al. (2013), remaining studies only provided indirect evidence of successful reproduction. Thus, further investigation in natural settings is needed in order to more broadly evaluate the possible impacts of C&R on reproduction.

Atlantic salmon (*Salmo salar*) is an emblematic species that supported subsistence fisheries for thousands of years and commercial as well as recreational fisheries for decades (MacCrimmon & Gots, 1979; Aas et al. 2010; ICES, 2019). Despite a generalised decline in abundance across its range, which led to closures and restrictions of fisheries (ICES, 2019, Atlantic salmon remains a highly prized game fish and is targeted by recreational anglers across the North Atlantic during its upstream spawning migration (ICES, 2019). In Eastern Canada alone, recreational fisheries targeting Atlantic salmon generates over $100 million annually which makes it a priority for conservation effort (ASF, 2011). Concerns about the specie’s conservation status led to the common adoption of C&R by fisheries managers (MFFP 2016, ICES 2019). The total number of C&R salmon have increased substantially over the years; in 2018, over 162 000 salmon have been released around the North Atlantic (ICES, 2019).

Across North America and Europe, implementation of captive-breeding programs is also used in combination with C&R to compensate decreasing recruitment on populations with abundance below their appropriate conservation threshold (DFO, 2008; MFFP, 2016; NASCO, 2017; ICES 2019). As for C&R, very few studies have documented patterns of reproductive success of captive-bred salmon (Milot et al. 2012; O’Sullivan et al. 2020). Previous research on captive-bred salmon demonstrated that minimizing time spent in captivity would alleviate unintentional domestication effect (Milot et al. 2012). Hence, there is a need to determine whether current hatchery practices still produce salmon with reduced reproductive success to assess the potential negative effect of those programs on targeted populations.

Global warming also necessitates involvement from Atlantic salmon managers to better understand the potential impact of elevated river temperature on released salmon. Indeed, previous studies have shown that the extent of post-release mortality varies with water tempetarure and to a lesser extent also according to angler practices, gear and bait types, angler experience (Cooke & Suski, 2005; Lennox et al. 2017). Although probability of post-release mortality is low (<0.05) at water temperature ranging from 0 to 12°C, it increases with warmer waters and expected to be as high as 0.35 at water temperatures ranging from 20 to 25°C (Van Leeuwen et al. 2020). Given the physiological impact of elevated river temperature on released salmon, evaluating its consequences on reproductive success is crucial to determine the long-term sustainability of C&R in the face of climate change (Gale et al. 2013).

Here, we evaluate relative reproductive success of captive-bred as well as C&R Atlantic salmon from a wild anadromous population spawning in the Rimouski River, Québec, Canada. This river has been targeted by several stocking programs since 1992. The current program is now gradually ending since the population recently reach its appropriate conservation target from 2017 to 2020 (MFFP, 2021). Captive-bred salmon are still present in this population, which allowed comparing their reproductive success to that of their wild counterparts. We further tested the contribution of captive-bred salmon to the population’s genetic diversity and compared its relative effect to the contribution of their wild counterparts, including mature male parr, which are young salmon that sexually mature at very small size, roughly 200 times less than that of anadromous males. These small males have previously been shown to significantly contribute to preserve genetic diversity in salmon populations (Johnstone et al. 2013; Perrier et al. 2014). Moreover, the exploitation of the Rimouski R. salmon population by recreational anglers as well as C&R practice have increased over the last decades such that an average of 100 salmons are now released annually (MFFP, 2020). The Rimouski R. is also one of the warmest rivers used by salmon in Quebec which frequently exceeds 20°C during fishing season (MFFP, rivtemp.ca/mffp_qc/). Thus, this system allows assessing the contribution of C&R salmon to the reproductive output of the population and their ability to reproduce when released at elevated water temperature that are expected to increase post-release physiological impacts and mortality.

Using novel microsatellite sequencing technology, we accurately quantified the reproductive success of captive-bred Atlantic salmon and C&R salmon by linking putative parents with their young-of-the-year progeny with molecular parentage analyses. The five goals of this study were to; i) test whether current captive-breeding programs produce Atlantic salmon with reduced reproductive success compared to their wild counterparts; ii) identify the possible factors responsible for the discrepancy in their reproductive success; iii) compare the contribution of captive-bred salmon and mature male parr to genetic diversity; iv) determine if C&R salmon have a reduced reproductive success when compared to non-caught salmon; v) evaluate the effect of temperature at release on reproductive success of C&R salmon.

## Material and Methods

### Study site

The Rimouski River is located on the South shore of the St. Lawrence Estuary, Québec, Canada (48°44′ N; 68°53′W). On average, 723 adult Atlantic salmon spawners returned to the river annually from 2014 to 2018 (MFFP, 2020). A dam acting as a complete barrier to upstream migration is located four km upstream to the river mouth, at the level of a natural impassable waterfall. During their upstream migration, adult fish are trapped in a cage and transported 1 km above the dam to be released upstream. Spawning grounds and suitable habitats for Atlantic salmon juveniles extend over a stretch starting from the dam and ending 21 km upstream at a second impassable waterfall. Spawning also occurs downstream of the dam, this minor section of the river and the salmon that remained there are therefore not included in this study.

### Captive-breeding program of Rimouski River

The captive-breeding program of Atlantic salmon in the Rimouski R. started in 1992 and from 1992 to 2019, a total of 716 176 fry, 884 624 parr and 297 019 smolts have been stocked. Starting in 2012, the broodstock was comprised of about 20 males and 20 females, with a semi-factorial crossing design using a minimum of 3 females and 3 males. One third of the broodstock held in captivity was replaced on an annual basis and no individual spawned more than three times. For this stocking program, the number of juveniles to be stocked annually (about 60 000) was determined based on a mathematical model described by Bernatchez (2009) in order to result in at least a 15% increase in abundance while limiting the effective population size reduction at less than 10%. Broodstock where collected from the Rimouski R. each year and crossed at the Québec Government hatchery located in Tadoussac. None of these captive breeders were born in the hatchery and kept all its life in captivity. However, some returning adult trapped at the dam and brought to the hatchery each year to maintain the pool of breeders may have been themselves offspring from captive-born fish that had been released as fry, parr or smolt in the Rimouski R. during past supportive breeding years, when stocked salmon where not marked. From 2012 to 2017, stocked salmon were young-of-the-year weighting 3.21 to 3.54 g. Adipose fins were removed from stocked salmon for future identification purposes.

### Atlantic salmon recreational fishery in the Rimouski River

Recreational Atlantic salmon angling on Rimouski R. is restricted to summer months beginning June 15^th^ and closing September 30^th^. Anglers have to register at the Rimouski’s ZEC (Area of Controlled Exploitation) which monitors recreational fishing activities on the river. Provincial regulation allows angler to keep up to two grilse (salmon < 63 cm in fork length) or to catch and release 3 salmon, whichever quota is reached first. All large salmon (> 63 cm) must be release.

### Sampling

#### Adult sampling and stocked fish identification

Measurements and fin samples were taken on all anadromous adults crossing the dam in 2018 when fish were released across the dam. The genetic sex of every returning adult was determined using King & Stevens (2019) PCR-amplification based method. Adult were identified as hatchery born using the presence/absence of adipose fin which was clipped on captive-bred fish.

#### Caught and released fish sampling

During the 2018 fishing season, angler participation was requested to collect tissue samples on their C&R salmon (punch of 5mm diameter of adipose fin) and to gather information on the C&R event. To facilitate the sampling process, anglers received punch pliers coupled with a kit that contained a 1.5 ml Eppendorf filled with 95% ethanol to preserve the adipose punch and a water-proofed sheet to record information on the C&R event. After each C&R event, anglers recorded date, time of the day, the pool number, the length of the fight (from hooking to landing), the kind of hook used (single or double, barb or barbless), the hooking location on the fish, the presence of bleeding and approximate air exposure duration. Temperature data loggers were placed in all frequently visited river pools. Given the information collected by the anglers, we could record the water temperature for each C&R event. Angler participation was not mandatory; however, the scientific team was present every fishing day on the main river pools to promote the project and eventually assist anglers during the sample collection.

#### Fry sampling

From July 15^th^ to August 15^th^ 2019, young-of-the-year (or fry, age 0+) born in the river were sampled using electrofishing over a stretch starting from the dam and ending 21 km upstream at an impassable waterfall. The very small tributaries of this river stretch do not provide known spawning grounds or significant habitat for juveniles because of their limited accessibility. Fry sampled were conserved in a 15 ml Falcon tubes filled with 95% ethanol. Within every 1 km reach, we selected 12 electrofishing sites (mean area 100 m^2^) given their fry habitat suitability index which was provided by the Québec Minister of Forests, Wildlife and Parks (MFFP). Each site was electro-fished once to maximise the amount of site sampled in a single day. To assess the effect of the number of offspring sampled on the potential for detection of anadromous parents, we sub-sampled 50 to 2381 offspring by steps of 50, with each step being subsampled 1000 times. In the case where enough offspring were sampled, the number of identified parents should reach a plateau.

### Molecular analyses

DNA was extracted from adult’s adipose fin tissue and fry’s caudal fin tissue using salt extraction method described by Aljanabi and Martinez (1997). Fifty-two microsatellite loci were then amplified by PCR in two multiplexes previously developed by Bradbury et al. (2018) (panel 1a and 2a). Locus amplification followed the protocol of Zhan et al. (2017); two sets of polymerase chain reactions (PCRs) were used, a multiplex PCR and an index PCR. Each multiplex PCR was performed using Qiagen Multiplex Master Mix. Each oligonucleotide in the multiplex reaction was tailed with Illumina sequencing primer sequence and served as oligo-binding sites in the subsequent index PCR. PCR multiplex conditions were the same for the two multiplexes and included 5 ul Qiagen Multiplex Master Mix, 10 ng DNA, 2.4 ul Oligo Mix for a total volume of 25 ul. Thermal reaction conditions included 95°C, 25X (94°C 30s, 65°C 3m, 72°C 30s) 72°C 30s. Multiplex-PCR products were pooled (per sample) in equal volume amounts, cleaned using Quanta-Bio SparQ Puremag Beads, and then used as template for the index PCR. Libraries were sequenced at 10–12 pM concentration at the genomic platform of the Institut de Biologie Intégrative et des Systèmes (IBIS), Université Laval, Québec (http://www.ibis.ulaval.ca/). Sequencing was performed using Illumina MiSeq (Illumina) and the MiSeq Reagent Kit V3 with 150 cycles in one direction and dual indexing. Indexed individuals were demultiplexed with the Miseq Sequence Analysis software. Allelic sizes were then scored using MEGASAT (Zhan et al. 2017) setting minimum depth (per sample per locus) at 20 reads; i.e. alleles with less than 20 reads were not called. Examination of histogram outputs (depth vs allele size) from MEGASAT confirmed allele scores, and we adjusted scores when necessary. Loci with more than 10% missing data were removed from the dataset to insure precision to our parentage analysis.

### Parental allocation of fry

Parentage allocation was conducted using Cervus v 3.0 (Kalinowski et al. 2007; Marshall et al. 1998) and Colony2 (Jones & Wang, 2010) that differ in their approach to parentage assignment. Cervus3 uses simulated parents and offspring to determine cut-off points of log-likelihood (LOD) scores for true parents, which are then used to identify parent-offspring pairs in empirical data. Cervus3 was first used to find the most probable mother-offspring and father-offspring dyad. As mentioned above, Rimouski R. is known for harboring mature male parr. Those males were not sampled in 2018; thus, the full likelihood approach implemented in Colony2 (Jones and Wang, 2010) was used to infer their genotypes from the pedigree analysis. Briefly, Colony2 uses a group-wise method to find the most likely configuration of full-sib and half-sib families in the data. Since we sampled every mother in our system, grouping offspring into full-sib and half-sib families allowed us to infer the genotype of mature male parr given the genotype of the mother when the offspring was not already assigned to an anadromous father.

### Detection of first generation (F_0_) migrant

Mobley et al. (2019) have shown that local Atlantic salmon exhibit higher fitness in their native river compared to dispersers. Therefore, we conducted a first generation (F_0_) migrant detection analysis with Geneclass2 (Piry et al. 2004) to remove those potential dispersers from downstream analysis. Since we had no genotypic reference of potential source populations with this microsatellite panel, we computed the likelihood of individual genotype within the Rimouski population (L_home). To do so, we used the frequency-based method of Paetkau et al. (1995) and the Monte Carlo resampling algorithm developed by Paetkau et al. (2004) simulating 10 000 individuals to estimate the probability of an observed multi-locus genotype. Individuals that had a probability below 0.01 (type I error) were considered as F_0_ migrant, and thus removed from downstream analyses.

### Analysis of relative reproductive success (RRS) of hatchery and C&R fish

We estimated relative fitness of hatchery-born fish and C&R fish to that of wild-born fish and uncaught fish, respectively. To do so, we used the number of offspring assigned to a given spawner as a measure of reproductive fitness. This measure is necessarily partial because not all offspring of a given spawner were collected. Then, we computed the ratio of reproductive fitness of the hatchery/wild fish and the C&R/non-caught fish which gives a measure of relative reproductive success (RRS) using Eq. 14 of Araki & Blouin (2005). RRS calculations were conducted independently for single-sea-winter (1SW) males, multi-sea-winter (MSW) males and females. Analyses were performed separately for each sex since they may respond differently to the hatchery environment and C&R events (Christie et al. 2014). To test for statistical significance, we perform non-parametric one-tailed permutations to test the hypothesis that captive-bred and catch-and released fish have lower fitness than that of wild and uncaught fish, respectively (i.e. whether RRS < 1.0). Briefly, numbers of offspring assigned to each parent were permutated

1 000 000 times (without replacement) and the probability of obtaining a value equal of larger than the observed fitness difference was evaluated (see Araki & Blouin 2005 for details). One-tailed test were chosen as we had an *a priori* expectation for lower RRS in captive-bred based on previous work.

Following Araki & Blouin (2005) approach, we used the maximum-likelihood method developed by Kalinowski & Taper (2005) to calculate confidence intervals for RRS estimates. Their method is appropriate for our study because it assumes that the frequencies of hatchery and wild fish are known exactly and that the only uncertainty in the estimate of RRS comes from sampling a finite number of offspring. Considering this set of assumptions, maximising the number of offspring sampled allowed us to obtain more precise estimates of the realized RRS regardless of the adult sample size.

Reduced RRS in captive-bred fish can generally be explained by reduced survival of their offspring in the wild because of transgenerational effects of the hatchery on offspring phenotypes via genetic or epigenetic inheritance. To rule out the possibility that hatchery fish could simply have lower spawning success or, in the case of females, a higher rates of egg retention, we tested whether a higher fraction of captive-bred fish would have a reproductive success equal to zero due to spawning failure. Thus, using the prop.test function implemented in R, we tested if there was a higher proportion of zero reproductive success in hatchery fish compared to wild fish.

### Effects of captive-breeding on components of reproductive success

Then, we assessed the captive-breeding effects on two components of reproductive fitness, namely the number of offspring assigned and the number of mates. The main idea was to evaluate which of the two components was significantly different between captive-bred and wild salmon. Results from this analysis would therefore help better understand significant deviation from one in RRS estimates. We performed this analysis separately for females and males using a zero-inflated negative binomial model (ZINB). We applied a ZINB model because estimates of number of offspring assigned and mating success contained a high proportion of individuals without any offspring, which hampers the possibility to identify their mates. Zero-inflated mixture models consisted of a binomial model for the frequency of zeros, and conditional on this, a count model using a negative binomial distribution. For females, we performed individual analysis per mate type, that is alternative reproductive tactics (mature male parr, 1SW and MSW males). For each model explaining the number of offspring assigned to female, we built a global ZINB model using origin (i.e. captive-bred/wild), fork length and number of mates. For the analysis pertaining to the number of mates, we built a global ZINB models using origin and fork length. To analyse the factors influencing the number of offspring assigned to males, we built a global ZINB model using origin, number of years spent at sea (one (1SW) or more (MSW)) and number of mates as well as the interaction between time at sea and origin and between time at sea and number of mates. For the analysis pertaining to the number of mates for males, we built a global ZINB models using origin, time spent at sea and number of mates and the interaction between time at sea and origin.

For all models, we assessed the fit of the global model by visualizing quantile-quantile residuals distribution and rootogram which graphically compares empirical frequencies with fitted frequencies from a given probability model with the package countreg in R (Zeileis & Kleiber, 2018). Then, we built a set of models made of all models nested within the global model (i.e. all combinations of including or excluding each variable) plus a null model (intercept only). Those models were ranked according to their AICc (a corrected measure of AIC for small samples), and ΔAICc (AICc of the model minus the AICc of the best model) was computed for each model. Following this procedure, we built a confidence set of models with all models with a ΔAICc < 4 (according to Hurvich & Tsai, 1989). Finally, we quantified the effect of variables appearing in the top models with multi-model inference using the shrinkage method (Burnham and Anderson 2002). Every model was build using the ‘pscl’ package in R 3.4.4.

### Genetic estimates of the effect of captive-bred fish on the effective number of breeders (Nb)

To investigate the contribution of captive-bred fish on the effective number of breeders (Nb), this parameter was estimated for three different datasets corresponding to parent-fry assignments implicating (i) wild anadromous breeders (*n* = 916), (ii) wild and captive-bred anadromous breeders (*n* = 1560) and iii) wild anadromous breeders and mature male parr (*n* = 1544). To do so, we used the LDNe program (Waples & Do 2008) to estimate Nb and the contribution of wild and captive-bred fish to Nb. We used a threshold of 0.05 as the lowest allele frequency that gives the least biased results according to Waples & Do (2010). Since the sample of parent-fry assignment to wild breeders only was smaller than that also including the contribution of mature male parr, we subsampled these latter datasets using the same number of fry as that of the wild breeders dataset. In this way, the threshold on the lowest allele frequency has the same effect on each dataset. Hence, we subsampled the parent-fry assignment of data sets ii) and iii) 1000 times and calculated Nb on each subsampling step. Finally, we compared value of Nb obtained in group i) with the distribution generated with group ii) and iii).

### Estimate of the effect of mature male parr and captive-bred salmon on offspring’s genetic diversity

To contrast the contribution of mature male parr to allelic richness to that of captive-bred males, we compared the total number of alleles found among progeny assigned to i) wild anadromous pairs, ii) wild and captive-bred anadromous pairs and iii) wild anadromous female and mature male parr for an increasing number of offspring sampled. We subsampled from 50 to 1500 fry, by steps of 50, 1000 times for each step, and estimated the total number of alleles found among fry assigned to the aforementioned groups of parents. We represented the differences in the number of alleles among these three groups with a Loess regression of the mean value of the total number of alleles found for each number of fry sampled considered, as well as the 95% distribution of values.

### Effect of temperature and air exposure time on reproductive success of C&R salmon

To assess the effect of the variable of the C&R event on the reproductive output of MSW adults, we used a negative binomial model since it describes count data that shows overdispersion. Because we had data for a limited number of C&R salmon only, we limited the number of predictor variables included in the model. Therefore, we tested the relationship between the effect of temperature during the C&R event, the air exposure time and their interaction on RS since they are variables known to affect RS of C&R fish (Richard et al. 2013). To do so, we built a global negative binomial model using the aforementioned variables with the pscl package in R (Jackman, 2020). To assess the fit of the global model, we visualized quantile-quantile residuals distribution and rootogram which graphically compares empirical frequencies with fitted frequencies from a given probability model with the package countreg in R (Zeileis & Kleiber, 2018). Then, we built a set of models made of all models nested within the global model plus a null model (intercept only). We ranked the best models using AICc (corrected measure of AIC for small samples), and we computed the model minus the ΔAICc of the best model). This step allowed to build a confidence set of ΔAICc < 4 (as suggested by Hurvich & Tsai, 1989). Finally, we quantified the effect of variables appearing in the top models with multi-model inference using the shrinkage method (Burnham & Anderson, 2002).

## Results

### Adult population of Rimouski River

From June 15th and October 30th of 2018, 475 anadromous Atlantic salmon (273 males and 202 females) were transported above the dam, of which 26 % were from the captive-breeding program (*n* = 126) (Table 1). The anadromous population was composed of 57 % (*n* = 271) salmon > 63 cm which are considered as MSW and of 43% (*n* = 204) salmon < 63 cm which were considered 1SW salmon. The proportion of MSW and 1SW was significantly different between wild and captive-bred salmon (*χ*^2^= 7.58, p-value = 0.005) with the former having 186 (53%) MSW for 160 (45%) 1SW and the latter having 85 (67%) MSW for 41 (33%) 1SW. Eight individuals were identified as first-generation migrant, of which two were MSW females and six were SSW males. In total, Colony 2 estimated that 432 mature male parr fathered 750 offspring with a mean reproductive success of 1.69 offspring per mature parr. The presence of mature parr highly skewed the sex ratio which was of 1 female to 1.43 male before accounting for them and of 1 female to 3.7 males when accounting for them.

**Table 1:** Details of the genotyped adults and juveniles Atlantic salmon. The samples include all the returning adults caught at the Rimouski River dam from summer 2018 and the juveniles caught on the Rimouski River spawning grounds during spring 2019.

|  | Number of individuals |
| --- | --- |
| Adult transported above dam | 475 |
| Adult used for assignments * | 468 |
| Born in the river (wild origin) | 349 |
| Born in the Rimouski hatchery | 126 |
| Returning as SSW | 204 |
| Returning as MSW | 271 |
| C&R fish | 33 |
| C&R fish transported above dam | 18 |
| Fry assigned | 2495 |
| * Difference between the number of adults used for assignment and the number transported above the dam is due to fish with missing or partial genotypes. |  |

### Caught-and-released salmon during 2018 angling season

A total of 33 fish out of 89 C&R salmon were successfully sampled by anglers. Most of those C&R events were done below the dam (31/33), three km upstream from the mouth of the river and one km downstream from the dam. Identity analysis between salmon that crossed the dam and C&R salmon revealed that 18 of those salmon successfully crossed the dam after being released (Table 1). Thus, we obtained a reproductive success measurement for a total of 13 MSW female (mean ± sd length = 78.1 ± 5.6 cm), 4 MSW male (length = 76.1 ± 4.79 cm) and 1 SSW male (length = 55 cm).

### Parental allocation

After filtering for loci with more than 10% missing data on both parents and offspring data set, 38 polymorphic microsatellite loci were retained for subsequent analyses. The number of alleles ranged from 3 to 13 per locus (average = 7) and observed heterozygosity ranged from 0.024 to 0.8396 with an average of 0.55. Average F_is_ was -0.005 (95**%** CI: -0.01-0.0091) thus indicating the absence of within-river Wahlund effect.

For the 475 adults transported above the dam, we obtained a complete genotype of 468 individuals that was then used for parental allocation (Table 1). A total of 2495 fry were sampled during July and August in 2019, genotyped and assigned to putative parents that spawned above the dam during fall 2018. We unambiguously assigned the paternity of 1617 and the maternity of 2495 offspring samples to anadromous adults and the paternity of 750 offspring to mature parr. The remaining 128 fry were assigned to mature parr with a probability lower than 0.8 and were thus excluded from subsequent analyses.

### Description of the mating system

The number of inferred offspring to females ranged from 0 to 67 (mean = 5.6, variance = 76.3) for mature male parr mates, from 0 to 29 (mean = 3.5, variance = 22.7) for 1SW mates and from 0 to 33 (mean = 4.4, variance = 33.5) for MSW mates. For males, the number of inferred offspring ranged from 1 to 8 (mean = 1.7, variance = 0.9) for mature male parr, from 0 to 23 (mean = 3.7, variance = 21.9) for 1SW salmon and from 0 to 56 (mean = 11.6, variance = 173) for MSW salmon.

Females were highly polyandrous with three females that mated with one male only. The total number of mates per female ranged from 0 to 33 (mean = 7, variance = 38.0). Mature male parr mates per female ranged from 0 to 26 (mean = 3.6, variance = 16.9), 1SW mates per female ranged from 0 to 7 (mean = 1.6, variance = 2.8), MSW mates per female ranged from 0 to 8 (mean = 1.6, variance = 3.0). Males were polygamous but to a lesser extent than females: 57 males mated with only one female of which 47 were 1SW males. The number of mates for mature male parr ranged from 1 to 5 (mean = 1.4, variance = 0.4), for 1SW it ranged from 0 to 7 (mean = 1.64, variance = 2.43) and for MSW it ranged from 0 to 16 (mean = 4.2, variance = 16.9).

### Relative reproductive success (RRS) of captive-bred versus wild and C&R versus non-caught Atlantic salmon

Estimates of RRS for captive-bred versus wild and C&R versus non-caught salmon are shown separately for males and females and time spent at sea (1SW, MSW) in Table 2. Our results show that captive-bred salmon had a reduced fitness compared to their wild counterparts. Thus, the RRS of MSW hatchery-reared females was of 0.805; this value was significantly lower one (p < 0.01) given our permutation test and our maximum-likelihood confidence interval estimation (Figure 1). For MSW hatchery-reared males, RRS was of 0.828; likewise, this value was significantly lower than one (p < 0.01) given both our permutation test and our maximum-likelihood confidence interval estimation (Figure 1). The RRS of captive-bred 1SW males was of 0.649 which again was significantly lower than one (p < 0.01) wild born 1SW males.

**Fig 1.**
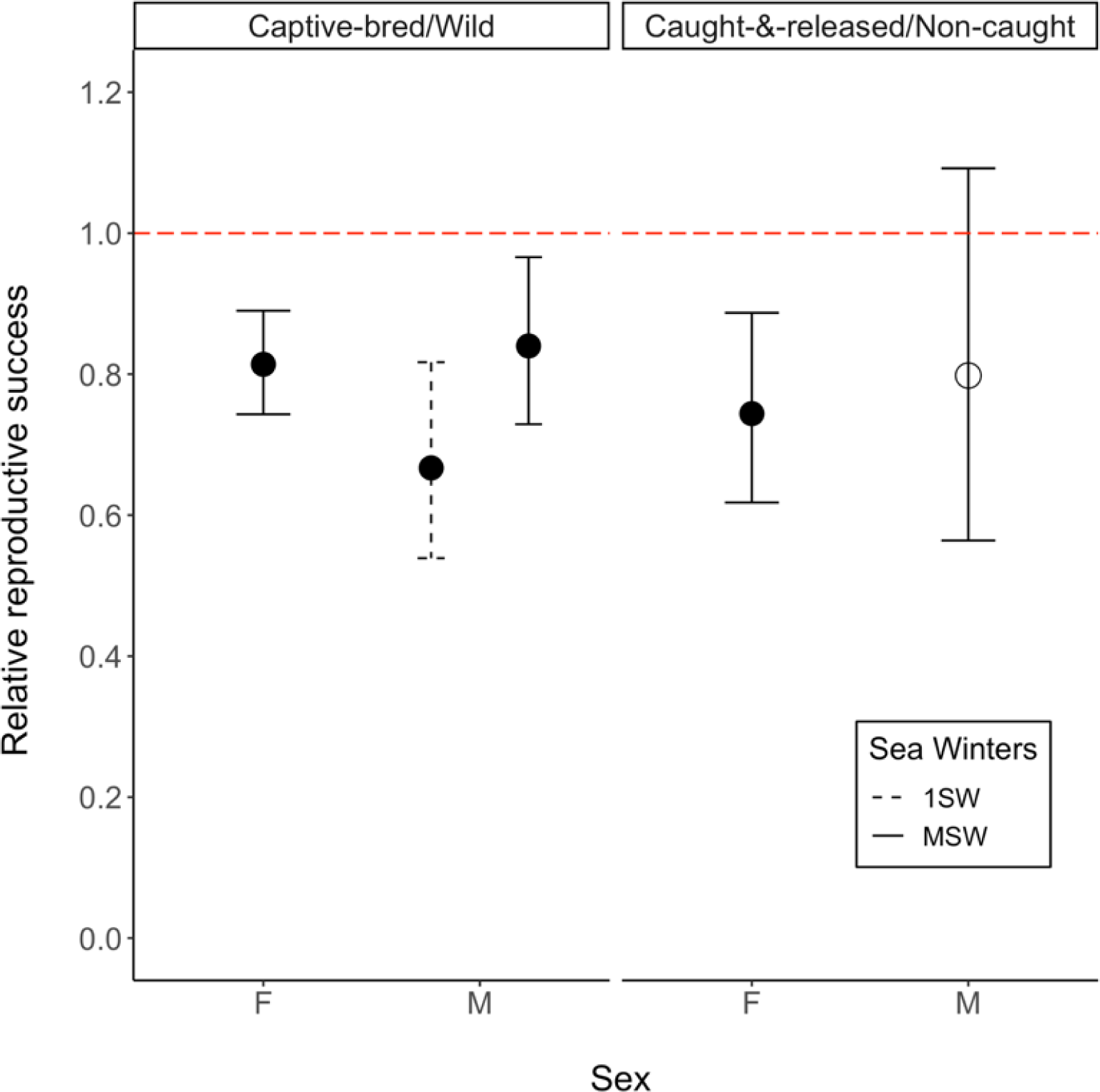
Maximum-likelihood estimates of relative reproductive success (RRS) and their associated 95% confidence intervals for captive-bred vs wild fish and caught-&-released vs non-caught Atlantic salmon. If captive and wild or C&R and non-caught salmon had equal fitness, then RRS would be equal to 1 (dashed red line). Filled and empty circles represent RRS that were significantly different at the 95% confidence interval given the permutation test.

**Table 2:** Relative reproductive success (RRS) of naturally spawning F_1_ parent. (*n*F1 is the sample size for naturally spawning captive-bred (C-B), wild (W), caught-and-released (C&R) and non-caught (N-C) parent; *n*F2 is the number of offspring assigned to each group of parents. RS is the reproductive success measured as the mean number of offspring assigned per parent. Variance is the average of the squared differences from the mean reproductive success. RRS is calculated as the RS of captive-bred fish over the RS of wild-origin fish and RS of C&R fish over the RS of non-caught fish, associated P-values are based on one-tailed permutation tests. Statistical power is the RRS value that would be significant with 80% and 95% probability).

| Sex/<br>Winter at<br>sea | $n F_1$<br>(C-B/W) | $n F_2$<br>(H/W) | RS<br>Captive-bred | Variance<br>Captive-bred | RS<br>Wild | Variance<br>Wild | RRS | P-value | 80%/95%<br>Power |
| --- | --- | --- | --- | --- | --- | --- | --- | --- | --- |
| Female<br>MSW | 53/133 | 607/1860 | 11.5 | 101 | 14.0 | 238 | 0.805 | <<br>0.01 | 0.963/0.929 |
| Male MSW | 28/51 | 286/620 | 10.2 | 160 | 12.2 | 183 | 0.828 | <<br>0.01 | 0.943/0.897 |
| Male 1SW | 40/146 | 105/593 | 2.62 | 11.8 | 4.06 | 24.3 | 0.649 | <<br>0.01 | 0.919/0.853 |
| | $n F_1$<br>(C&R/<br>N-C) | $n F_2$<br>C&R/N-C) | RS<br>C&R | Variance<br>C&R | RS<br>Wild | Variance<br>Wild | RRS | P-value | 80%/95%<br>Power |
| Female<br>MSW | 13/174 | 124/2362 | 9.54 | 77.8 | 13.5 | 206 | 0.732 | <<br>0.01 | 0.932/0.868 |
| Male MSW | 4/75 | 37/869 | 9.25 | 236 | 11.6 | 173 | 0.797 | 0.07<br>41 | 0.754/0.866 |

For C&R fish, C&R MSW females had a RRS of 0.732 which was significantly lower (p< 0.01) than one given our permutation test and our maximum-likelihood confidence interval estimation. C&R MSW males had a RRS of 0.797 but this value was not significantly different than one given both our permutation test and maximum-likelihood confidence interval estimation (Figure 1). Hence, our results show that C&R tend to have a reduce fitness compared to non-caught salmon, but this reduction was only significant for females.

### Effect of origin and male reproductive tactics on components of reproductive success

The number of offspring assigned to females increased with the number of mates for all three male alternative reproductive tactics (Table 3, Figure 2, A)). However, as expected, mating with mature male parr increased the number of offspring assigned at a lower rate than for 1SW and MSW mates. Female length had a significant positive effect on the number of offspring assigned when mating with MSW males, however its effect size was negligeable compared to the effect of number of MSW mates. The number of offspring assigned to wild and captive-bred females did not differ significantly, suggesting that the survival rate of offspring from captive-bred females was comparable to that of wild females. Females’ mating success showed a positive relationship with length for all three types of male alternative reproductive tactics (Figure 2, B)). There was no difference between the number of mature male parr and MSW mates between captive-bred females and wild females. However, captive-bred females had significantly less 1SW mates than wild females. Therefore, the observed reduction of RRS of captive-bred females was apparently mainly caused by a lower mating success with 1SW mates compared to wild females.

**Fig 2.**
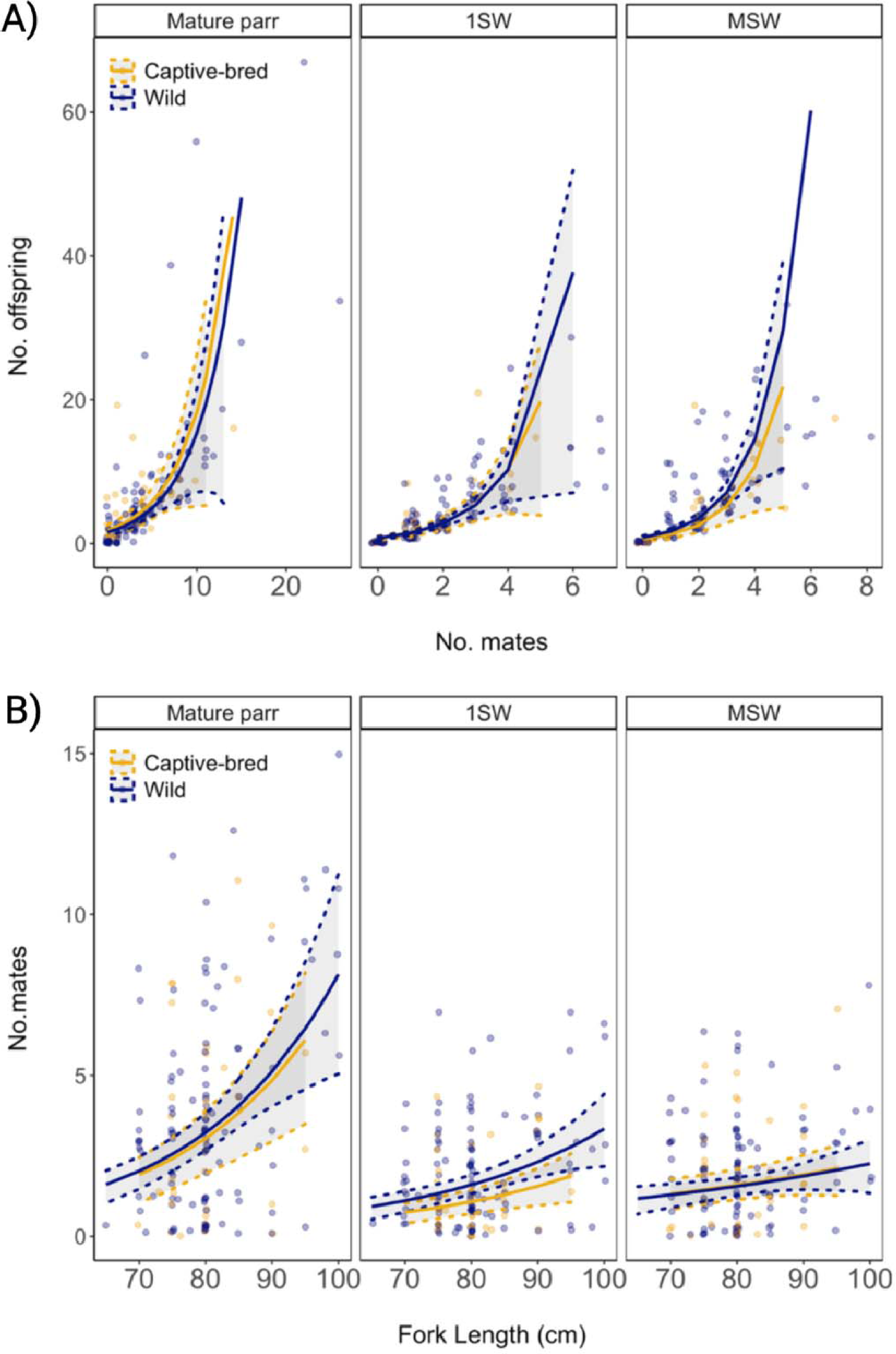
A) Relationship between the number of offspring (No. offspring) and mating success (No. mates) for female Atlantic salmon for three alternative reproductive tactics (Mature parr, 1SW and MSW). Coloured lines represent ZINB model prediction for captive-bred (blue) and wild (yellow) females, grey areas and hatched lines represent 95% confidence intervals (CI) obtained by bootstrap. Circles show individual data points. B) Relationship between mating success (No. mates) and length of female Atlantic salmon for three alternative reproductive tactics (Mature parr, 1SW and MSW). Coloured lines represent ZINB model prediction for captive-bred (blue) and wild (yellow) females, grey areas and hatched lines represent 95% confidence intervals (CI) obtained by bootstrap. Circles show individual data points.

**Table 3:**
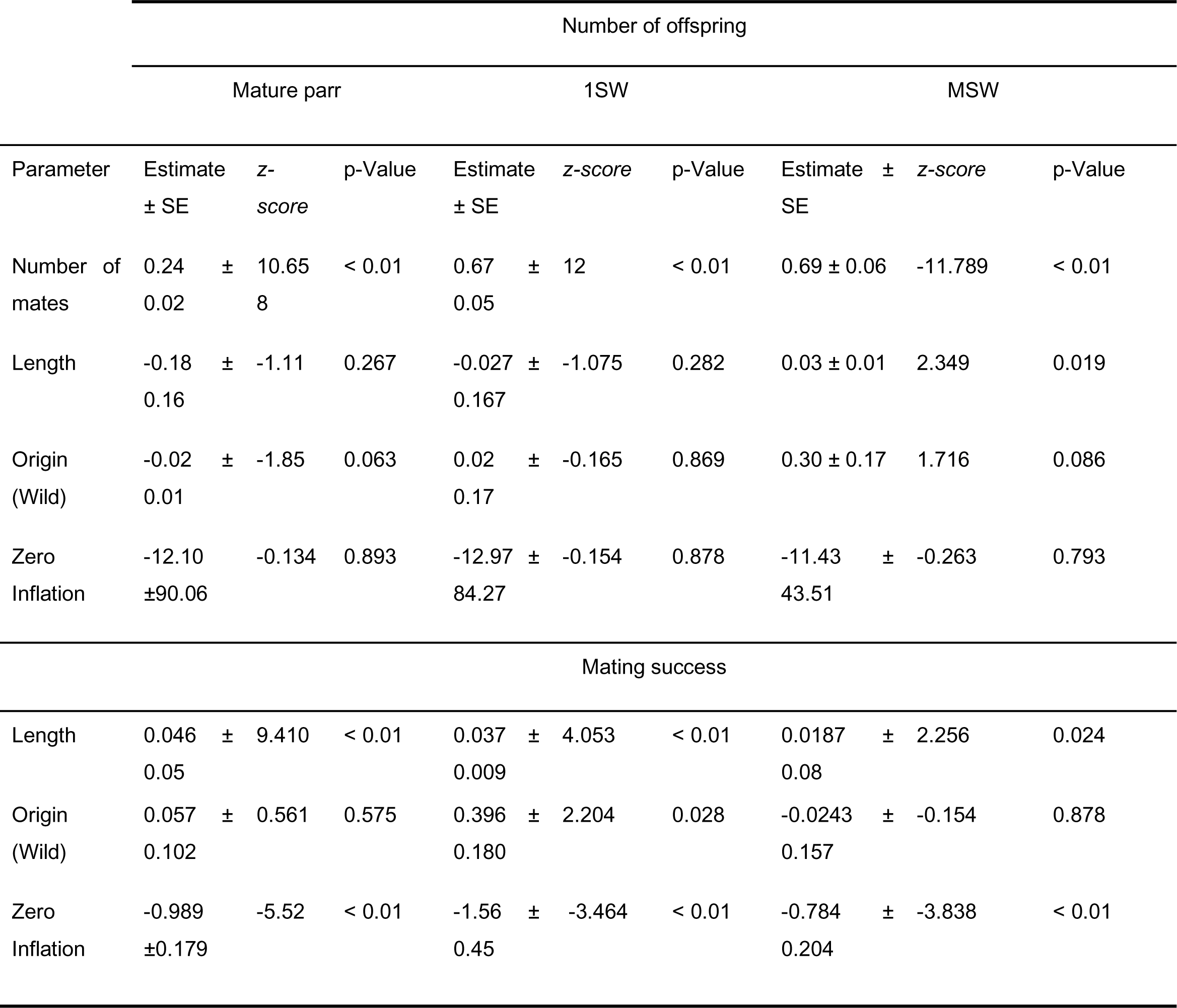
Summaries of ZINB model testing the effect of number of mates, length and origin (wild/captive-bred) and the effect of length and origin on the number of offspring assigned to females and mating success respectively. The “zero inflation” term accounts for the large number of adults with zero reproductive success and mating success in our sample.

MSW males had a higher number of offspring assigned than the 1SW salmon (Table 4, Figure 3, A)). Moreover, both 1SW and MSW males increased their number of offspring by mating with multiple females but 1SW males did so more rapidly than MSW males. As for captive-bred females, captive-bred males did not show a significant reduction in offspring assigned relative to their wild counterparts. 1SW males had fewer mating partners than MSW males and both 1SW and MSW captive-bred males showed a significant reduction in their mating success compared to their wild counterpart (Figure 3, B)). Hence, the previously observed reduction of RRS for both 1SW and MSW captive-bred males was apparently mainly caused by a lower mating success compared to wild males.

**Table 4:** Summaries of ZINB model testing the effect of number of mates, sea age, and origin (wild/captive-bred) and the effect of sea age and origin on the number of offspring and mating success respectively. The “zero inflation” term accounts for the large number of adults with zero reproductive success and mating success in our sample.

| Parameter | Number of offspring |  |  |
| --- | --- | --- | --- |
| | Estimate ± SE | z-score | p-Value |
| Intercept | 0.74 ± 0.18 | 4.23 | < 0.01 |
| Number of mates | 0.28 ± 0.03 | 10.59 | < 0.01 |
| Sea age (1SW) | -0.90 ± 0.21 | -4.37 | < 0.01 |
| Sea age (1SW) : Number of mates | 0.36 ± 0.06 | 6.18 | < 0.01 |
| Origin (Wild) | -0.01 ± 0.13 | -0.08 | 0.93 |
| Zero inflation | -12.27 ± 78.35 | -0.157 | 0.876 |
| Parameter | Mating success |  |  |
| | Estimate ± SE | z-score | p-Value |
| Intercept | 1.44 ± 0.13 | 10.97 | < 0.01 |
| Sea age (SSW) | -1.06 ± 0.12 | -8.67 | < 0.01 |
| Origin (Wild) | 0.28 ± 0.14 | 2.08 | 0.038 |
| Zero inflation | -1.90 ± 0.33 | -5.79 | < 0.01 |

**Fig 3.**
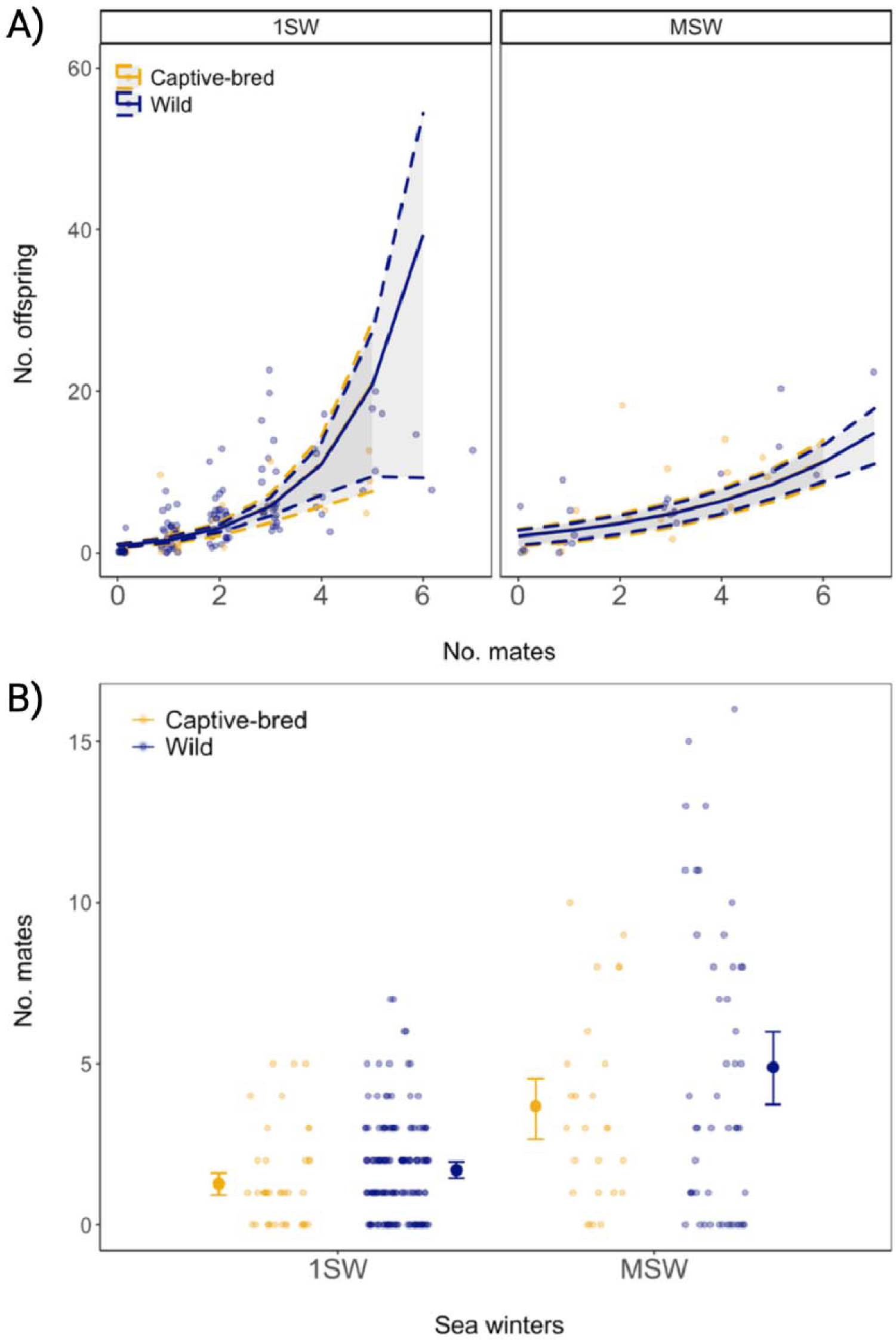
A) Relationship between number of offspring (No. offspring) and mating success (No. mates) for 1SW and MSW male Atlantic salmon. Lines represent ZINB model prediction for captive-bred (blue) and wild (yellow) 1SW and MSW males, grey areas represent 95% confidence intervals (CI) obtained by bootstrap. B) Relationship between mating success (No. mates) and winters at sea (Sea winters)x in male Atlantic salmon. Large circles with error bars represent the model prediction ± 95% confidence interval (CI) obtained by bootstrap while small circles show individual data points. Captive-bred males had significantly less mates than wild males (p < 0.038).

### Estimation of the effect of captive-bred anadromous salmon and of mature male parr on genetic diversity of offspring

Using LDNe to estimate Nb from the genetic data, we obtained values of 114 (CI: 110.8-121.7) for wild anadromous salmon only, of 146 (CI: 136.8-154.9) for wild and captive-bred anadromous salmon combined and of 173 (CI: 160.5-185.6) for all wild anadromous salmon and mature male parr (Figure 4). Thus, captive-bred salmon increased the effective number of breeders by 28% which is significantly less compared to mature male parr which inflated it by 52%. Global estimate of Nb obtained from COLONY, including wild, captive bred and mature male parr was of 271 (CI: 231-323).

**Fig 4.**
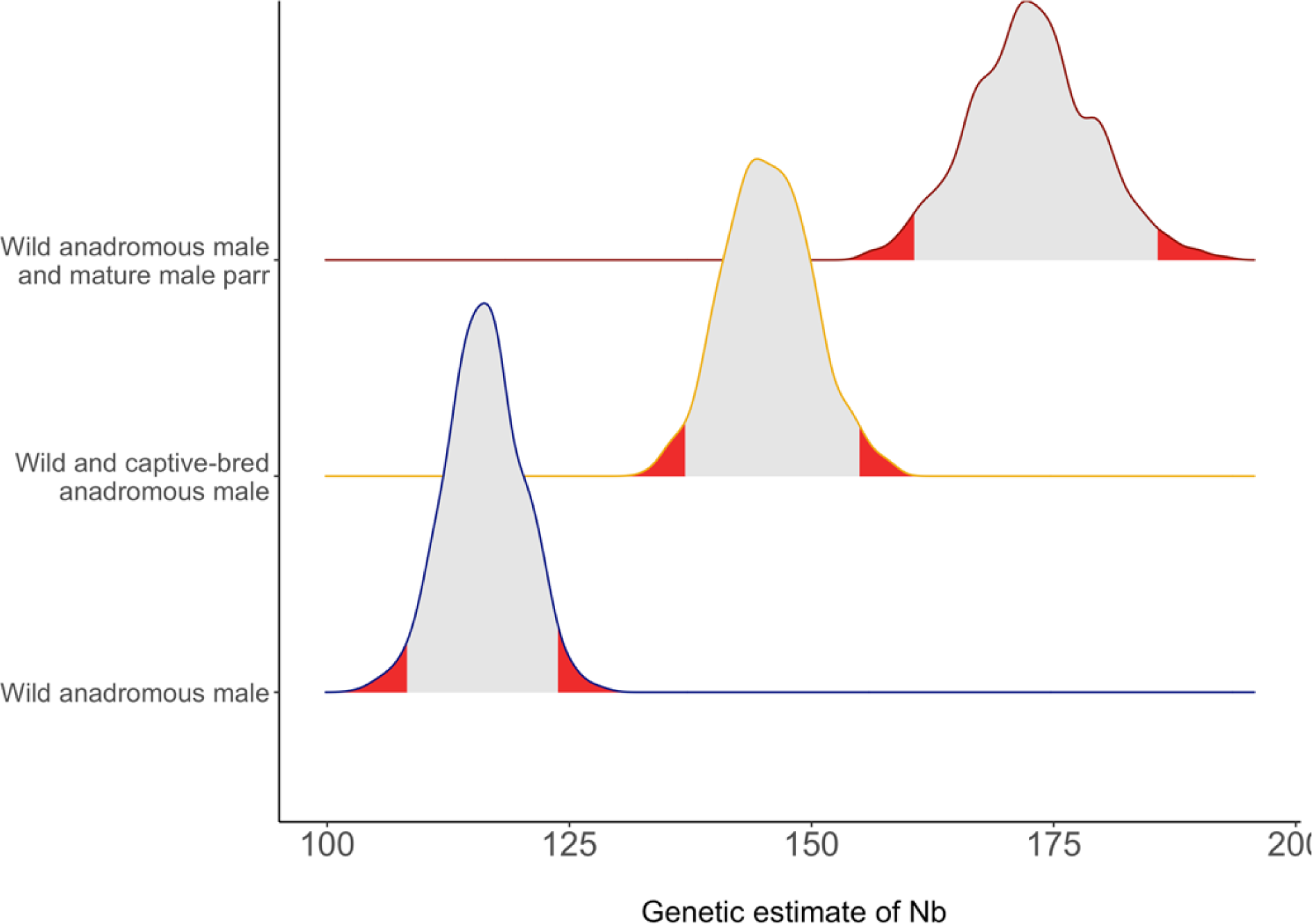
Distribution of genetic (LDNe) estimates of effective number of breeders (Nb) considering either only wild anadromous salmon, both wild and captive-bred anadromous salmon or both wild anadromous salmon and mature male parr. The distribution of Nb calculated for wild anadromous salmon only was obtained directly from NeEstimator 2 (Do et al. 2014) whereas the distribution for wild and captive-bred anadromous salmon or wild anadromous salmon and mature male parr was obtained by subsampling 916 fry from the parent-fry assignment of wild and hatchery breeders 1000 times and calculating Nb on each subsampling step. The 2.5% and 97.5% tails of the distribution are indicated in red.

Captive-bred anadromous salmon and mature male parr both contributed to increase the total number of alleles in the population relative to considering wild anadromous fish only (Figure 5). Fry produced by pairs of anadromous females and mature male parr had a higher number of alleles than fry produced by pairs of wild anadromous salmon and by wild and captive-bred anadromous salmon. For instance, sampling 1000 offspring 1000 times in each dataset, the average number of alleles was 252 (CI: 252-259) for wild anadromous salmon, 258 (CI: 254-262) for wild and captive bred anadromous salmon and 265 (CI: 265-272) for wild anadromous salmon and mature male parr.

**Fig 5.**
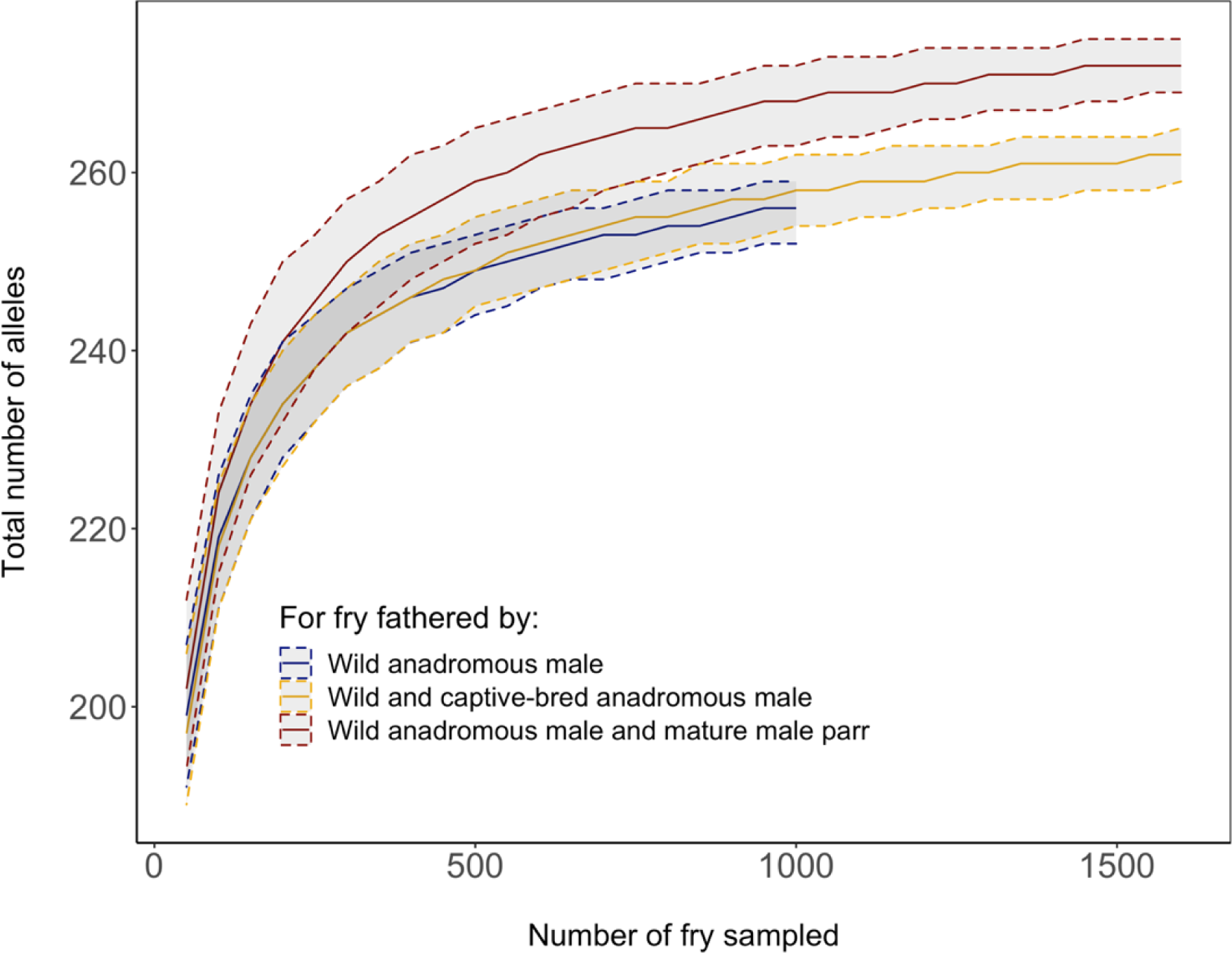
Loess regression of the mean value of the number of alleles over all genotyped microsatellite loci calculated for 50 to 1500 fry fathered by wild anadromous male, wild and captive-bred anadromous male combined and all wild anadromous male and mature male parr. Each estimate was bootstraped 1000 times. Five to 95% interval distribution of the data were given around the mean value.

### Impact of C&R conditions on reproductive success

The best model was the null model which provided the lowest AICc whereas the global model had the highest value. This result indicates that neither air exposure nor water temperature had a significant effect on the number of produced offspring. The effect of temperature on reproductive success of C&R salmon is shown on Figure 6. Surprisingly, the C&R event that occurred at the higher water temperature was associated with the highest observed reproductive success among fish manipulated by anglers. These results must be interpreted cautiously given the small sample size that was available for this part of our study.

**Fig 6.**
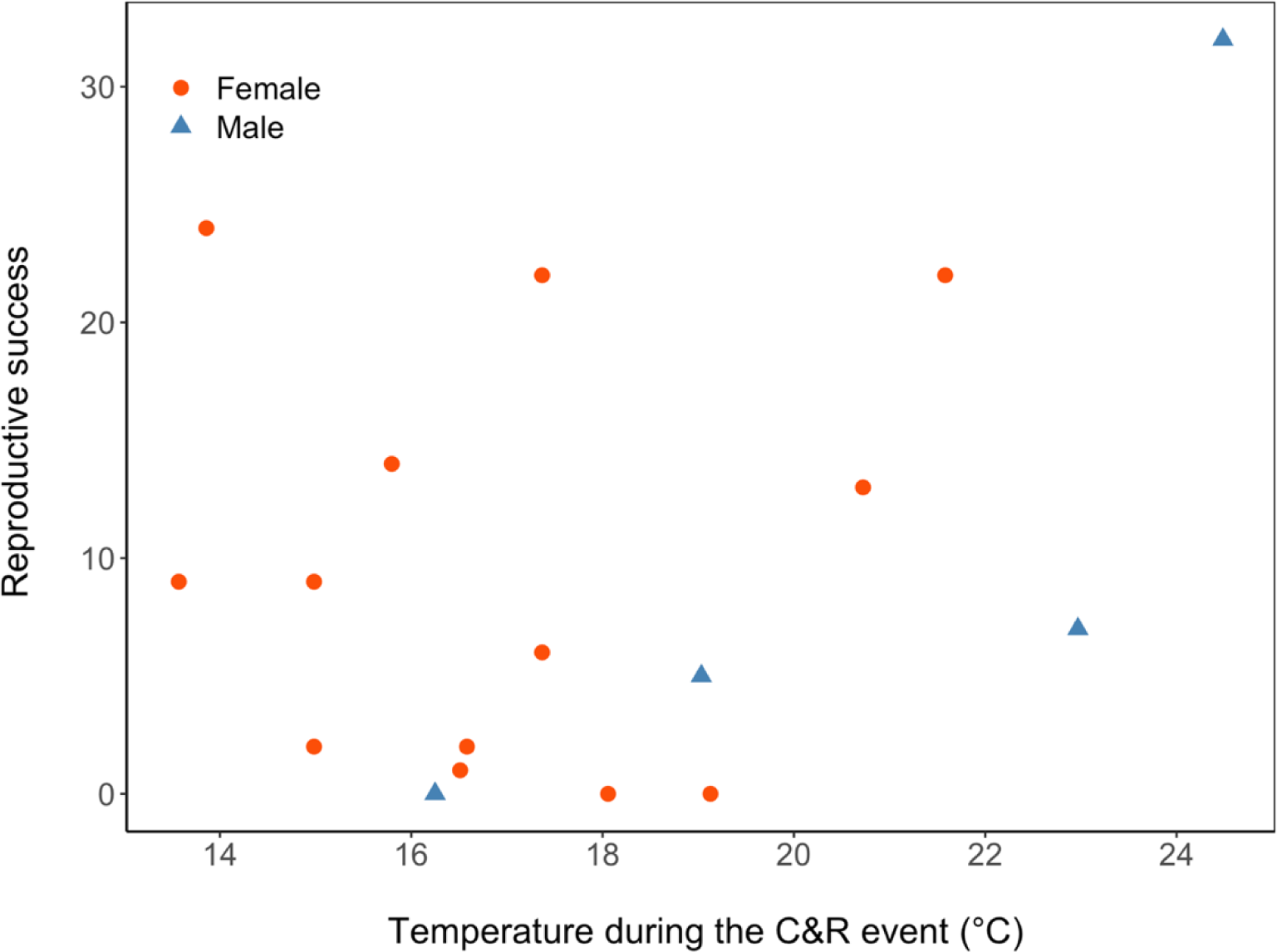
Reproductive success (estimated by number of fry assigned) for the group of caught-and-released Atlantic salmon (n = 18) with increasing water temperature.

## Discussion

Most wild populations of Atlantic salmon have experienced a sharp decline in abundance over the past century. As a consequence, captive-breeding programs for fish supplementation and adoption of catch-and-release in recreational fisheries have generally been implemented as a mean to preserve ecosystem integrity, enhance declining populations, and sustain fisheries. In this study, we evaluated the efficiency of these practices by means of molecular parental assignment in a wild population of Atlantic salmon from the Rimouski River, Québec. We demonstrate that captive-bred males and females displayed lower RRS than their wild counterparts mainly because captive-bred salmon had a lower mating success. Yet, captive-bred salmon contributed to increase genetic the effective number of breeders in the wild population, albeit to a lesser extent than mature male parr which also contributed to increase allelic diversity. Finally, we provide evidence that caught-and-released males and females have reduced RRS compared to salmon that have not been caught, but we found no evidence that elevated water temperature has an effect on their reproductive output. Below we underline the evolutionary and conservation implications of these results which should contribute to guide future directions for management of Atlantic salmon populations.

### Atlantic salmon mating system in the Rimouski River

#### Female reproductive and mating success

Female reproductive success was highly variable and best explained by their number of mates. All females mated with multiple anadromous males and mature parr which reflects the well documented male-biased operational sex-ratio in this species (Fleming et al. 1996). In such conditions, display of aggressive behavior by MSW males has a low reproductive pay-off because the high relative number of male competitors decrease the probability of successfully defend spawning females (Fleming & Gross, 1994). Hence, multiple male spawning event occurs, and sperm deposition is usually in order of dominance status (Fleming et al. 1996, 1997). Accordingly, female reproductive success increased more rapidly for MSW mates than for 1SW and mature parr mates which adopt a sneaking strategy (Fleming & Einum, 2011).

Interestingly, length was also a significant, albeit weak predictor of female reproductive success when mating with MSW males. Potential egg production of female closely tracks increase in body size (Fleming, 1996); yet several studies also found a weak effect of female length on their reproductive success (Taggart et al. 2001; Garant et al. 2001, 2003; Richard et al. 2013). This is not completely unexpected since environmental heterogeneity in time and space plays an important role in determining offspring survival (Gibson & Myers, 1988). Rivers draining on the south shore of the St. Lawrence River watershed are generally characterized by lower annual minimum flows which is a known abiotic factor that reduce survival of juvenile salmon (Assani et al. 2006). Low river discharge is associated with a reduction in subsurface flow, and therefore oxygen supply in the gravel substrate, which in turns have negative effects on developing eggs and embryos (Gibson & Myers, 1988). Such stochasticity in abiotic factor is expected to decrease the relationship between body size and reproductive success among females (Chapman, 1988).

Polyandry has been reported in several studies on Atlantic salmon and our study confirms that it is the main reproductive strategy in the Rimouski R. Such mating pattern has direct genetic benefits for female; for example, Garant et al. (2005) found that females with a higher number of mates also had more outbred offspring, and that both of these characteristics increased their reproductive success. Moreover, in environmentally heterogeneous systems, such as the Rimouski R., polyandry could also have potential ecological benefits. Female Atlantic salmon can reduce variance of fitness by adopting multiple-redds strategy depending on temporal and spatial variation in offspring survival (Barlaup et al. 1994). Admittedly, without behavioural observation, we cannot tell whether multiple matings occurred at different nest sites rather than at a specific redd. Nonetheless, we found a spatial distribution of families that is highly heterogeneous with half-siblings from the same mother frequently collected more than 5 km apart (Bouchard, pers. obs.). These observations have also been reported in Garant et al. (2001) and strongly suggests that females might have constructed multiple redds since fry generally have low dispersal ability (< 1.5 km, Eisenhauer et al. 2020).

#### Male reproductive and mating success

As for females, there was high interindividual variance in reproductive success among males which was best explained both by number of mates and the number of years spent at sea. This is in accordance with the fact that Atlantic salmon males do not provide any parental care, making their offspring production an increasing function of mating success (see Arnold & Duvall, 1994). However, our results showed that the relationship between mating and reproductive success differs among alternative reproductive tactics. Although MSW males are typically dominant on the spawning territories and achieve higher reproductive and mating success than 1SW males (Garant et al. 2003), they exhibit courtship behavior, actively fight and defend nesting site against other males which in turn, increase the cost of mating with multiple females (Fleming, 1996). On the other hand, 1SW males are much smaller than MSW salmon, and behave like subordinate males on the spawning grounds (Fleming, 1998). Instead of fighting, these males have to cruise between multiple females to increase their chances of successful mating. Our results thus suggest that achieved reproductive success of MSW males may represent a trade-off between investing into courtship and active defense of nests and the benefits gained from mating with multiple females. In contrast, 1SW increased their fitness more rapidly with multiple matings which reflects the benefits of their reproductive tactics when operational sex-ratio is highly male biased.

#### Mature male parr

We identified 432 mature parr male which fathered 30% of all offspring sampled. This is in broad accordance with previous studies which reported proportions varying between 22 and 65% (Garcia-Vazquez et al. 2001; Taggart et al. 2001; Saura et al. 2008; Weir et al. 2010). Compared to a previous study performed in Les Escoumins R. draining on the North shore of the St. Lawrence River in Québec, mean reproductive success of mature parr was lower (1.7 vs 2.4; Richard et al. 2013). Because the aggressive behaviour of anadromous males can restrain the participation of mature parr in reproduction (Jones & Hutchings, 2001; Tentelier et al. 2016), it seems plausible that the spawning success of male parr was reduced by the larger proportion of anadromous males in our study (57% vs 41% in Richard et al. 2013).

#### Reduced fitness of captive-bred Atlantic salmon after one generation of captivity

The reduced RRS of captive-bred Atlantic salmon corroborates results from Christie et al. (2014) who quantitatively synthesised the results of five studies that used genetic parentage analysis to estimate the fitness of first-generation hatchery-born adults and wild-born adults spawning in the wild in various salmonid species. In 46 out of 51 estimates, adults born in a hatchery had lower fitness than wild-born adults. However, compared to the one previous study on Atlantic salmon (Milot et al. 2013), captive-bred salmon from the Rimouski R. had a higher RRS. Here, captive-bred MSW females and males averaged ∼80% while 1SW males averaged 65% of the reproductive success of their wild counterparts compared to Milot et al. (2013) which reported an average relative reproductive success of 50% for both sexes. Furthermore, our ZINB model showed that captive-breeding did not directly affect the number of offspring per mating event but instead the number of mating events. Thus, the number of offspring per captive-bred females was not significantly reduced when controlling for mating success and length which suggests they were able to find suitable spawning sites and display level of fecundity similar than that of wild females. However, captive-bred females had fewer 1SW mates than their wild counterparts. Fleming & Gross (1993) previously reported that captive-bred females were less competitive in their ability to acquire a nesting territory during direct competition with wild females which constrained them to build fewer nests (Fleming & Petersson, 2001). If captive-bred females did build fewer nests than wild females, MSW males could more readily defend their redds which would decrease their probability of getting fertilized by 1SW males. Again, behavioral observation would be required to confirm this hypothesis.

As observed for females, the number of offspring assigned to captive-bred males did not differ from that of their wild counterparts when controlling for mating success and time spent at sea, suggesting that their ability to fertilize female’s eggs is similar to that of wild males. Yet, both MSW and 1SW captive-bred males showed a significantly reduced mating success which affected 1SW to a greater extent. Reduced aggressiveness of captive-bred males has previously been documented in Atlantic salmon (Fleming et al. 1996). However, if captive-bred males displayed reduced aggression during spawning season, we would expect MSW salmon to be more affected since they are actively competing for access to females. A more plausible hypothesis could be that captive-bred males display more aggression than their wild counterparts. A previous study on captive-bred Atlantic salmon reported that progeny resulting from captive-breeding practices to be more aggressive than wild-born conspecifics (Blanchet et al. 2008) and that this behavioral syndrome can hold years after being released in nature (Fleming et al. 1997). Male and female salmon invest more than 50% of their total energy to reproduction, but 90% of the male loss is in somatic investments (Jonsson, 1991; Hendry & Beall, 2004). During spawning, combat between MSW is an energetically costly behavior that may terminate quickly with a bite after rushing at an opponent or intensify into a fight that results in several wounds (Jones, 1959). Engaging in too many prolonged combats could be detrimental for males as they can be reproductively active for about two months (Webb and Hawkins, 1989, Fleming et al. 1996). Being parsimonious with engaging in fight is therefore crucial for MSW males, and it can most probably pay to be careful and exhibit subordinate postures when meeting superior competitors (Fleming, 1996; Fleming et al. 1996). This would be particularly true for 1SW males which have better chances to furtively gain access to females rather than engaging into combat. Accordingly, reduced mating success affected their RRS to a greater extent compared to MSW males.

Although we do not have behavioral data to support this hypothesis, a previous study on Atlantic salmon controlling for the genetic background revealed that the environmental effects of hatchery-rearing up to the smolt stage (juvenile migrating at sea) could be significant on male behavior (Fleming et al. 1997). Whereas captive-bred males had similar levels of aggression, they were involved in more prolonged aggressive encounters and incurred greater wounding and mortality than wild males. As a consequence, those males were less able to monopolize spawning and obtained a mating success 51% that of wild males. Reduction of mating success was less striking in our study, but captive-bred salmon were stocked in the Rimouski R. at the fry stage which significantly reduced exposure to the hatchery environment and may have contributed to increased RRS compared to salmon stocked at the smolt stage (Milot et al. 2013).

Early environment is a deterministic factor that acts on later performance in fish (Jonsson & Jonsson, 2014). Although differences in fitness between hatchery-reared and wild salmon were previously thought to be genetically based only, mounting evidence also points toward an epigenetic basis associated with the effect of hatchery environment on epigenetic reprogramming of the progeny produced in captivity (Christie et al. 2016; Le Luyer et al. 2017; Gavery et al. 2018; Wellband et al. 2021; Leitwein et al. 2021). Of particular interest is the study of Rodriguez Barreto et al. (2019) which showed that early-life hatchery exposure changed the pattern of methylation of Atlantic salmon males and that corticotropin-releasing factor receptor 1-like was one of the differentially methylated regions in hatchery fish. This gene is involved in increased aggression and activity in another salmonid, the Arctic charr, *Salvelinus arcticus* (Backström et al. 2015). Results from their study also demonstrated that those changes were inheritable which raises concern about the potential negative long-term consequences of captive-breeding. In our case, hatchery-rearing clearly reduced mating success of both females and males and further study should investigate whether a causal relationship exists between epigenetic modifications and aggressiveness of captive-bred Atlantic salmon.

#### Contribution of captive-bred salmon and mature male parr to the effective number of breeders (Nb) and their influence on genetic diversity

It is generally accepted that genetic diversity both within and between populations is important to conserve, and it is relevant to ask whether captive breeding programs effectively maintain genetic diversity (Fraser, 2008). In this study, captive-bred salmon significantly contributed to a 1.28-fold increase in the effective number of breeders. These results are in line with those or Araki et al. (2007b) which demonstrated that the use of local broodstock for stocking contributed to a 1.73-fold increase in Nb over two generations. The contribution of captive-bred salmon to increase Nb is encouraging from a conservation perspective since contemporary captive-breeding programs for Atlantic salmon generally aim to help population retain 90% of their genetic diversity over 100 years by maintaining a Nb of at least 100 (Waples, 1990). Although mature male parr fathered 30% of the fry that were genotyped, their contribution to reproduction led to a 1.52-fold increase in Nb. This is in broad agreement with a similar study performed in another salmon population from the Malbaie R. on the North shore of the St. Lawrence River where mature male parr resulted in a 1.79-fold increase in Nb compared to a Nb estimate without considering their contribution to reproduction (Perrier et al. 2014). Moreover, we demonstrate that the number of alleles found in pairs of wild anadromous female salmon and mature male parr was higher than that of wild and captive-bred anadromous salmon. Hence, mature male parr most likely buffer against reduction of genetic diversity in populations with relatively small census size and as such, could contribute to enhance their evolutionary potential and ultimately, the long-term persistence of fisheries.

### Effect of catch-and-release on the reproductive success of Atlantic salmon

#### Reduced reproductive success of released salmon

We found that the catch-and-release events reduced the reproductive success of female and male salmon by 28% and 20% respectively, although the reduction was significant for females only. Although released males did not have a significantly reduced reproductive success compared to uncaught males, there is a possibility that the qualitative difference was real, but that the statistical power of our analysis was limited by the small sample size for this part of the study (Christie et al. 2012).

The mechanism causing the reduced reproductive success of C&R salmon could be manifold. C&R event has various physiological consequence for fish (Gale et al. 2013). From hooking to landing, levels of extracellular acidosis increase conjointly with blood and muscle lactate causing a decreasing content of extracellular pH, plasma bicarbonate, ATP and glycogen which considerably reduce likelihood of recovery following release (Wilkie et al. 1996; 1997; Thorstad et al. 2003; reviewed in Gale et al. 2013). Coupled to exhaustive exercise is the production of “stress hormones” which result in a suite of physiological and biochemical alterations to the internal physiology of fish to maintain performance during exercise. This causes a shift of investment of energy from anabolic process (i.e., growth and reproduction) to catabolic activities (i.e., energy mobilization and restoration of homeostasis) (Bonga, 1997). This physiological cascade can lead to mortality in some cases but when it does not, it can result in so-called sub-lethal consequences (Schreck 2010). For instance, C&R can lead to disrupted gametogenesis because of reallocation of energy during reproductive maturation in sockeye salmon (Patterson et al. 2004) and to altered courting and mating behavior in smallmouth bass (Cooke et al. 2002). Given this, it is difficult to pinpoint the exact cause of the reduction of fitness of released salmon. Previous studies convincingly demonstrated that C&R does not affect gamete or fry quality in Atlantic salmon (Davidson et al. 1994; Booth et al. 1995). Nonetheless, recent simulation-based study on Atlantic salmon documented shortened migration distances in caught salmon relative to their non-caught counterparts, which likely arose from the stress and exhaustion experience after release (Lennox et al. 2016; but see Thorstad et al. 2003). This suggests that this prolonged stress and exhaustion could influence breeding success since it is highly linked to physiological condition on spawning grounds (Hendry and Beall, 2004). Ultimately, if hindered migration after release reflects reduced activity during spawning season, there could be fewer reproductive encounters or nests built by released salmon which would decrease overall fitness.

#### *Effect of temperature on reproductive success of released* salmon

This is the first study attempting to document reproductive success of released salmon above their optimal temperature for aerobic scope which is estimated at ∼20°C in Eastern Canada (DFO, 2012). Admittedly, our results must be interpreted cautiously given the small sample size that was available for this part of our study. Yet, our results suggest that increasing temperature did not affect the reproductive output of released fish. High temperature reduces dissolved oxygen concentration in water, while increasing the metabolic rate and thus the oxygen demand of fish. Ultimately, when oxygen consumption becomes less than oxygen demand, the fish physiology uniquely relies on anaerobic energy pathway. When the C&R event is paired with high water temperatures their impacts are synergistic, and the complete exhaustion of aerobic and anaerobic muscular fuels are possible (Wilkie et al. 1996). Consequently, survival of released salmon is highly dependent on water temperature and probability of mortalities can be as low as 5% at cool river temperature (<12°C) but range from 7% and 33% between 18 and 20°C (Lenox et al. 2018; Van Leeuwen et al. 2020). As mortality increases with water temperature, we would expect sublethal effects on released fish to increase as well. For instance, Richard et al. (2013) reported that increasing temperature negatively impacted reproductive success of salmon between 12-19°C. The fact that we find no such correlation might be related to the availability of thermal refugia (i.e. colder body of water) in our system. During high temperature events, Atlantic salmon often engage in behavioral thermoregulation by moving within available thermal refugia to alleviate physiological stress (Breau et al. 2011). All the released salmon in our study were caught close to a known thermal refugium which probably gave them the opportunity to escape high water temperature and to enhance restorative processes.

Another non-exclusive hypothesis is that this could be locally adapted to high water temperatures. Rimouski river frequently warm up to temperature higher than the Atlantic salmon lethal limit (>28°C) during practically every summer while sustaining a viable salmon population that support an angling fishery. Local adaptation for higher temperature tolerance has previously been demonstrated in other salmonids by shifting the temperature at which maximum heart rate ceases to increase (Eliason et al. 2011; Antilla et al. 2014). This would increase the amount of anerobic energy available to face C&R at high water temperature; therefore, increasing survival and minimizing sub-lethal effects.

#### Evolutionary and practical consequences of captive-breeding and catch & release on wild populations

This study provides a comprehensive overview of the potential benefits and consequences of captive-breeding and catch-and-release as management practices and raises issues pertaining to the conservation of wild Atlantic salmon populations. First, our results revealed that mating success was the main factor explaining the reduction in relative reproductive success of first-generation captive-bred salmon. A growing body of literature suggests that breeding in captivity generates epigenetic modifications in captive-bred salmon which may underly observed decrease of fitness of captive-bred fish. If such epigenetic modifications are inheritable, as suggested in a recent study (Leitwein et al. 2021) captive-breeding could have a cumulative negative effect on the reproductive success of spawners via introgressive hybridization of hatchery and wild stocks (Willoughby & Christie 2019). Further research should investigate whether epigenetic modifications truly underly the decrease in breeding success in Atlantic salmon. Meanwhile, we suggest that implementation of captive-breeding programs should be limited to the short-term (<10 years; Willoughby & Christie, 2019) when possible.

Second, we also showed that mature male parr contribute importantly to maintain genetic diversity. Anglers and managers generally pay little attention to the potential importance of mature male parr, either from a demographic or genetic standpoint. In fact, their presence has even been seen as having a negative effect on recruitment to fishery and on harvestable biomass (Myers, 1984). However, several studies now support their key role in maintaining the long-term genetic diversity within salmon populations (Araki et al. 2007; Perrier et al. 2014; Johnstone et al. 2013) and to reduce inbreeding (Perrier et al. 2014). Indeed, those males constitute a reservoir that can compensate for variations in the number of anadromous males returning from sea migration. This phenomenon, referred to as genetic compensation, is responsible for limiting the decrease of Ne when few anadromous male breeders survive until breeding (Araki et al. 2007b). Although we demonstrate that captive-breeding programs contributed to increase Nb in the Rimouski R. salmon population, it does so to a lesser extent than the contribution of mature male parr. Therefore, we suggest that the contribution of mature male parr to genetic diversity should always be considered and evaluated when managing Atlantic salmon populations.

Third, despite the fact that released salmon experienced reduced reproductive success (RRS) relative to that of their wild counterparts, it is noteworthy that a minimum of 83% of them successfully reproduced. These results corroborate studies showing that mandatory C&R resulted in a 2.3-fold increase in the number of spawning redds in Norway (Thorstad et al. 2003) and in higher parr and fry densities in Russia (Whoriskey et al. 2000). Moreover, we demonstrate that salmon released at river temperature above 20°C are able to successfully reproduce suggesting that those salmon were able to find thermal refugia or that the studied population is locally adapted to high water temperature. From a management and conservation perspective, these results provide evidence that mandatory C&R successfully increase sustainability of exploited fish population since released fish contribute to the reproductive output of the population and that this benefit might hold in the face of global warming. Raw data used for this study is available at the Dryad digital repository: to be completed after manuscript is accepted for publication.

## Acknowledgments

This study was funded by the Ministère des Forêts, de la Faune et des Parcs du Québec (MFFP) and the Canadian Research Chair in genomics and conservation of aquatic resources. R.B. was supported by scholarship from Fonds de Recherche du Québec - Nature et Technologies. We thank Stéphane Forest, William Cayer-Blais, Arnaud Benoit-Pépin, Félix-Antoine Picard, Juliette Bherer, Jerome Doucet-Caron and Cécilia Hernandez for field and laboratory assistance. Finally, we thank employees from ZEC Rimouski, Gilles Schooner from BORALEX and fishermen of the Rimouski River without whom this project would have been impossible. This project is part of the research program of Ressources Aquatiques Québec (RAQ).

